# Endophilin Recruitment via GPCR Interactions Enables Membrane Curvature Generation in the Absence of Anionic Lipids

**DOI:** 10.1101/2020.07.11.198937

**Authors:** Samsuzzoha Mondal, Imania Powers, Karthik Narayan, Samuel Botterbusch, Tobias Baumgart

## Abstract

The Bin/Amphiphysin/Rvs (BAR) family protein endophilin plays key roles in membrane curvature generation during endocytosis of cellular receptors. The Src homology 3 (SH3) domain of endophilin interacts with the proline rich third intracellular loop (TIL) of various G-protein coupled receptors (GPCRs). While electrostatic interactions between BAR domain and anionic membrane lipids have been considered to be the major driving force in curvature generation, it is unclear how the direct interaction between TIL and SH3 affects this function and its coupling with receptor internalization. Here we show that TIL mediated interactions alone not only recruit endophilin to the membrane but also facilitate curvature sorting and curvature generating behavior of endophilin. To demonstrate this, we designed model membranes with covalently lipid-conjugated TIL and lipids without net negative charge so that endophilin was recruited exclusively via SH3/TIL interactions. We find curvature generation and curvature sorting under those conditions. Furthermore, we show that TIL interacts electrostatically with membranes in the presence of anionic lipids and that this interaction can interfere with binding of SH3. Overall, our study suggests that an interplay between TIL, charged membranes, BAR domain, and SH3 domain mediate membrane curvature generation to regulate receptor endocytosis following receptor stimulation.

## Introduction

Endocytosis is an important regulatory pathway for G-protein coupled receptor (GPCR) mediated cell signaling processes (1, 2). Activated receptors can be downregulated through endocytosis and then either undergo lysosomal degradation or recycling back to the plasma membrane. This process modulates the number of available receptors on the plasma membrane (3, 4). During receptor endocytosis, multiple protein-protein and lipid-protein interactions at the membrane promote recruitment of endocytic accessory proteins and membrane invagination, followed by membrane scission (5–7). Understanding the orchestration of these interactions would help in identifying potential therapeutic targets to rectify misregulations of the receptors leading to pathological conditions (8).

The BIN/Amphiphysin/Rvs (BAR) superfamily protein endophilin is one of the key effectors in clathrin mediated endocytosis (CME), and particularly in clathrin-independent endocytosis (CIE) (9, 10). The dimeric protein on the one hand facilitates membrane remodeling the via crescent shaped BAR-domain dimer (referred to as N-BAR domain because of the presence of an N-terminal amphipathic helix). On the other hand, endophilin’s Src homology 3 (SH3) domain recruits other endocytic proteins such as synaptojanin (11) and dynamin (12), and interacts with several other proteins that contain proline-rich-domains (PRDs) (13). The multifunctionality of endophilin has given rise to major research interest into its role in CIE pathways such as fast endophilin mediated endocytosis (FEME). Endophilin plays a central role in FEME by driving cargo recruitment, membrane curvature generation, and membrane scission (14, 15). It has been suggested that molecular interactions occur between the endophilin SH3 domain and the third intracellular loop (TIL) of several GPCR family members during their internalization through the FEME pathway (10, 16, 17). Still, it has remained unclear how membrane remodeling by endophilin is functionally coupled with protein-protein interactions mediated by the SH3 domain.

Endophilin’s ability to sense and generate membrane curvature has been studied extensively *in vitro* as well as *in vivo*. Full-length endophilin and even the N-BAR domain alone can tubulate membranes when recruited via electrostatic interaction in the presence of anionic phospholipids (18–23). While binding to TIL of GPCRs provides an additional modality of membrane recruitment, its effect on membrane curvature generation has not been explored. In order to address this question, we specifically chose the TIL of the β1-adrenergic receptor (β1-AR) as it has been shown to bind the endophilin SH3 domain (14, 16). Interestingly, activation of β1-AR is found to trigger the FEME pathway in Retinal Pigmented Epithelial cells whereas β2-AR, another GPCR from the AR family whose TIL does not bind to endophilin, does not trigger FEME (14).

To study the effect of TIL-endophilin interaction on membranes *in vitro*, we have created a model membrane system by covalently coupling a TIL peptide from β1-AR to the lipid bilayer. This approach bypasses reconstitution of the entire GPCR and allows for precise control of the surface density of TIL peptides on the membrane. Earlier studies on curvature generation properties of endophilin used membranes composed of anionic phospholipids to facilitate recruitment of the protein to the membrane via electrostatic interactions with the N-BAR domain. While these electrostatic interactions have been considered to be crucial for curvature generations to what extent short range, non-electrostatic protein-lipid interactions contribute has not yet been explored.

To answer this question, we chose a neutral lipid composition to suppress electrostatic interactions of the membrane with either TIL or endophilin so that the membrane recruitment of endophilin could take place exclusively via covalently membrane-coupled TIL. We first show that TIL-mediated interactions can recruit endophilin to the membrane in the absence of any negatively charged phospholipids, and that these interactions could be interrupted by competitive electrostatic interactions between TIL and membranes containing anionic lipids. Our results reveal that endophilin retains its curvature sensing and generation properties even on uncharged membranes, which has far reaching implications for our understanding of how endophilin’s BAR domains interact with the membrane and are involved in receptor internalization processes.

## Results

### Covalent conjugation of TIL to lipid bilayers with zero net charge

This project aimed to investigate the consequences of endophilin’s interaction with the β1-AR GPCR in a lipid model membrane system. Reconstitution of a full length GPCR has been accomplished previously (24–26); however, receptor reconstitution would not allow us to control the number density of protein on the lipid membrane. Further complexity would be introduced since the GPCR-reconstitution process typically involves the use of detergents that interact with lipid bilayers and modify their properties (27). To circumvent these challenges, we covalently conjugated a known endophilin interaction domain, the receptor’s TIL (Figure 1A), with lipid headgroups. The TIL contains an intrinsic cysteine residue (Cys261 in β1-AR) located near the N-terminus of the TIL that allows formation of a covalent bond with a maleimide derivatized phosphatidylethanolamine lipid headgroup. We hypothesized that such a covalent conjugation would attach TIL on the membrane in a way that would enable the proline-rich domain to interact with endophilin’s SH3 domain.

**Figure 1.**
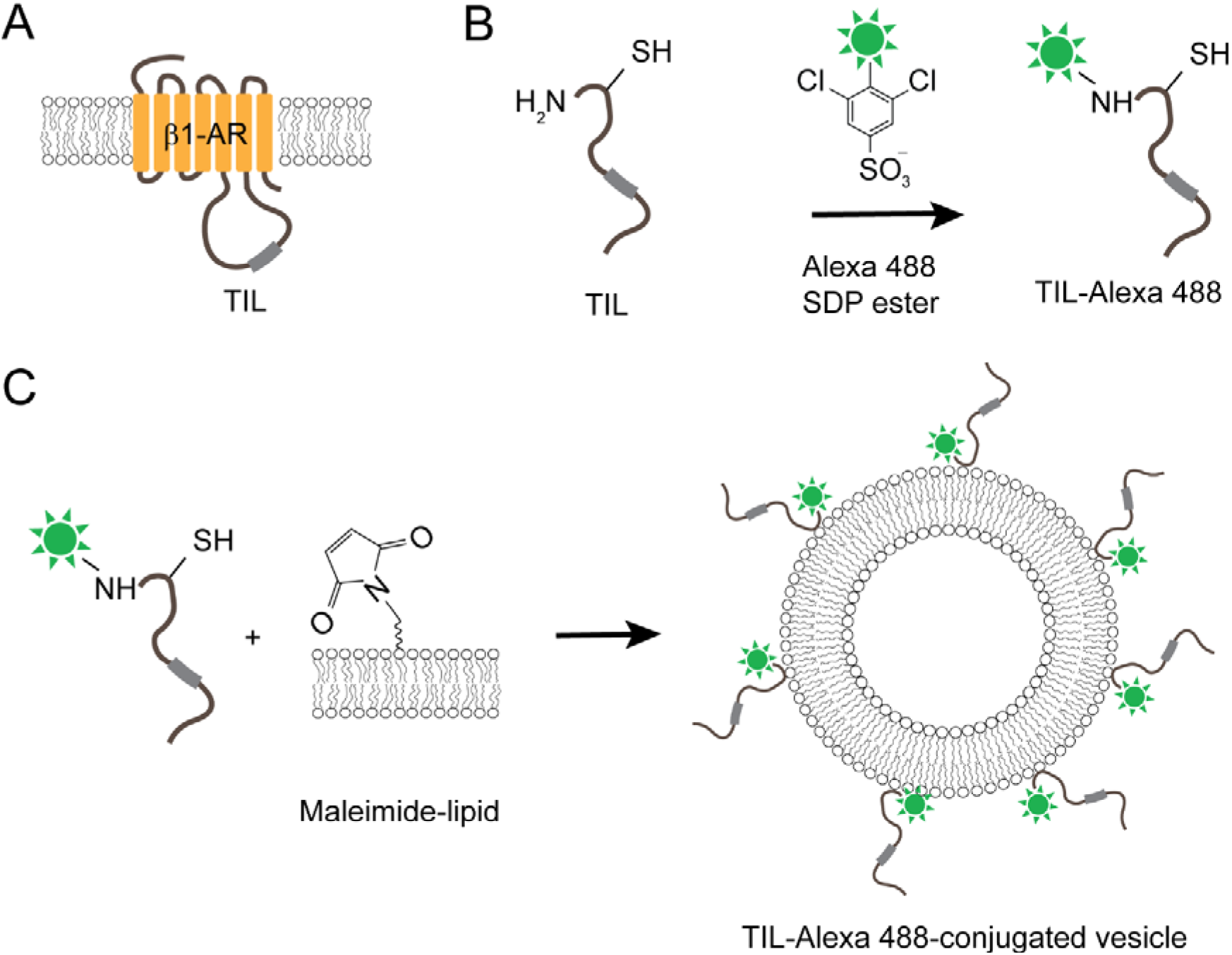
Design of model membrane system by covalently linking a GPCR TIL to the lipid bilayer. A. Cartoon representation of adrenergic receptor and position of TIL. B. Labeling of TIL peptide with Alexa 488-SDP ester allows selective modification of the N-terminus, leaving the cysteine side-chain free. C. Covalent conjugation of fluorescently labeled TIL to lipid bilayer containing lipids with maleimide-functionalized headgroup via cysteine-maleimide coupling reaction.

The purified TIL has a molecular weight of 8861 Da and is mostly disordered in nature (*Supplementary Information*, Figure S1 and S2). Using an orthogonal chemical reaction strategy, we attached the fluorophore to the N-terminal amine group of TIL and the fluorescently labeled TIL to maleimide-lipid. A commercially available SDP ester derivative of the fluorophore reacts with the N-terminal amine group leaving the cysteine–SH group free (Figure 1B). The reaction was performed at pH 7.0 because at this pH amine reactive probes react with the N-terminal amine group selectively over lysine sidechains (28). The molecular weight of the fluorophore conjugated TIL was confirmed by mass spectrometry (*Supplementary Information*, Figure S3). Further, incubation of the peptide with GUVs containing 5 mol% mCC-PE resulted in the coupling of the peptide on the membrane surface via cysteine-maleimide chemistry (Figure 1C). We verified this attachment on the GUVs by observing Alexa 488 fluorescence from the GUV surface via confocal microscopy imaging (Figure 2A).

**Figure 2.**
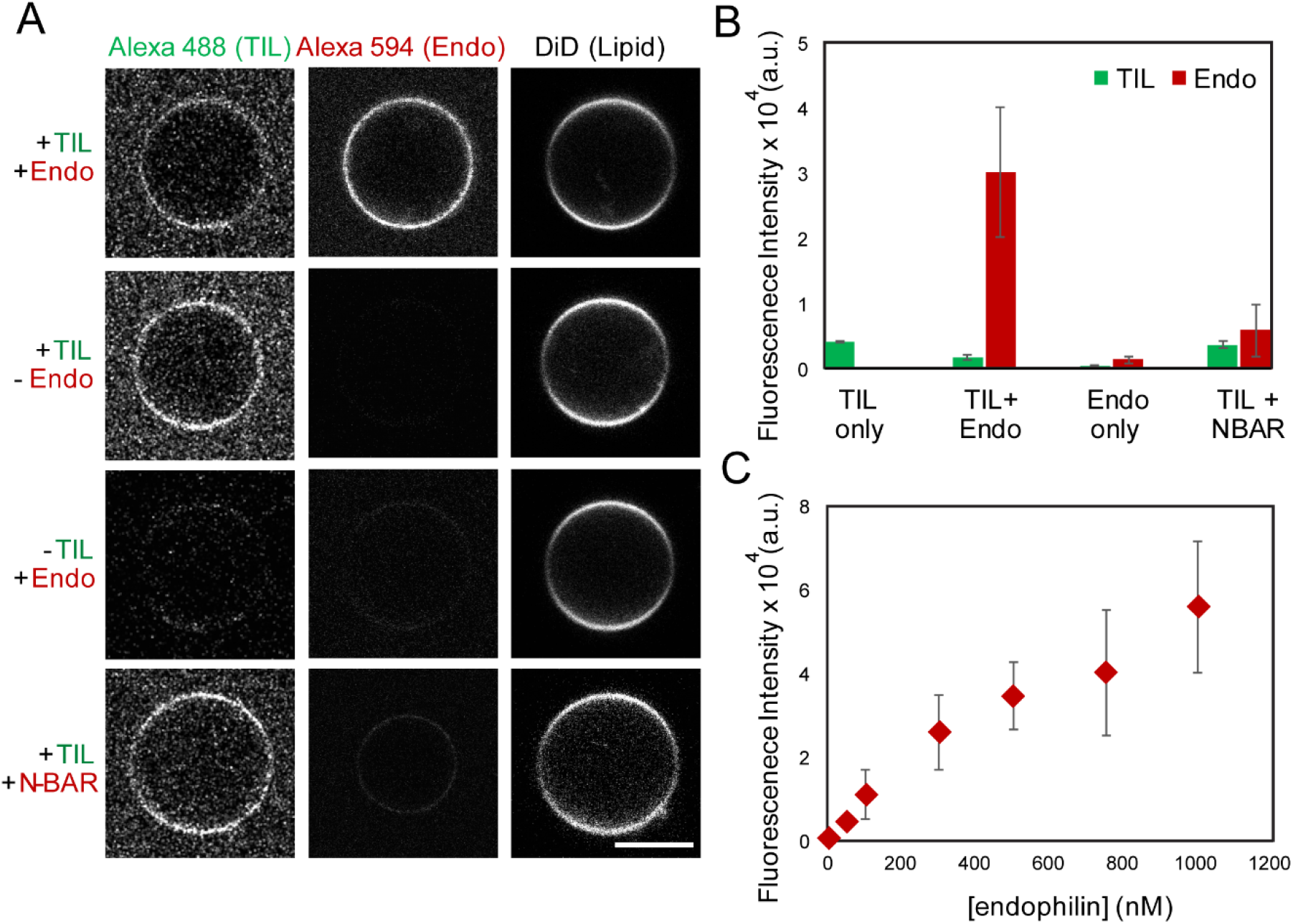
Endophilin can be recruited to membranes in the absence of anionic lipids, through membrane-bound TIL. A. Confocal images showing fluorescence channels corresponding to labeled TIL, endophilin (Endo) or its N-BAR, and of the membrane (through the DiD fluorophore). First row: demonstration of Alexa 594 labeled endophilin A1 binding to TIL-Alexa 488 conjugated vesicles (+TIL/+Endo). Second row: TIL-Alexa 488 conjugated GUVs in absence of endophilin (+TIL/-Endo). Third row: in the case of GUVs not conjugated to TIL (−TIL/+Endo), no significant recruitment of endophilin to the membrane is observed, as expected. Last row: unlike full length endophilin, it’s N-BAR domain is not recruited to the TIL-conjugated membrane (+TIL/+NBAR). All GUVs shown are composed of 94.5 mol% of DOPC, 5 mol% of mCC-PE, and 0.5 mol% of DiD. Scale bar 5 µm. B. Bar plot showing fluorescence intensities from the GUV images. C. A titration plot showing the fluorescence response from the TIL-conjugated GUVs in the Alexa 594 channel (Endo) as a function of endophilin concentration. Note that panel C suggests an apparent K_d_ value that is significantly higher than what was obtained in Ref. (16), which is expected since in addition to being recruited to the membrane, endophilin interacts with excess TIL in the solution. Data in the plots are represented as mean ± standard deviation (s.d.), N ≥ 3.

### TIL-SH3 interaction mediates recruitment of endophilin to lipid bilayers with zero net-charge

Earlier studies have found that endophilin A1 binds to the TIL of β1-AR with a dissociation constant of 47 nM (16). This interaction has been hypothesized to be a key element of the trafficking of this GPCR upon activation. Consequentially, we asked if this strong binding interaction would lead to recruitment of endophilin to our TIL-conjugated membrane surface even in the absence of any anionic phospholipids. A single cysteine mutant (E241C, C108S, C294S, C295S) of rat endophilin A1 was labeled with Alexa 594-maleimide, in order to visualize the protein on GUVs through confocal microscopy imaging. When we mixed the TIL-conjugated GUVs with Alexa 594-labeled endophilin and imaged within 10 minutes of incubation, we did observe binding of endophilin to the GUVs. This observation suggested that TIL recruited endophilin onto the membrane having no net negative charge (Figure 2A). A control experiment with GUVs of the same lipid composition but not conjugated with TIL showed negligible binding of the endophilin (Figure 2A, B). In order to confirm that this binding is mediated by SH3-TIL interaction only, we further tested an N-BAR only mutant of endophilin (since it would lack the SH3 domain). The N-BAR mutant shows significantly less fluorescence intensity from the TIL-conjugated GUV surface again suggesting that binding is mediated by the SH3 domain. The GUV imaging data confirm that endophilin can be recruited onto membranes by TIL even in the absence of anionic phospholipids.

### TIL binds electrostatically to negatively charged membranes

The TIL of β1-AR contains several lysine and arginine resides which result in a highly positive net-charge, with pI ~12. Such highly cationic segments are likely to interact with the anionic lipid headgroups within the inner leaflet of the plasma membrane. In order to test this hypothesis, we studied the binding of TIL with giant unilamellar vesicles (GUVs) consisting of a lipid mixture designed to mimic the negatively charged inner leaflet of the plasma membrane (PS/PE/PC 45:30:25). When we imaged the GUVs in the presence of Alexa 488-labeled TIL through confocal fluorescence microscopy, TIL showed binding to the GUVs (Figure 3A). To investigate if this mode of binding is primarily electrostatic or not, TIL and GUVs were mixed under various ionic strength conditions generated by changing the NaCl content of the mixing buffer in the range of 0-200 mM (Figure 3A, B). A steady increase in the extent of binding with a decrease in the ionic strength confirmed that the major mode of the TIL-lipid bilayer interaction is electrostatic.

From the ionic strength dependent binding studies, we observed that TIL shows a weak but significant membrane interaction at physiological salt concentration (~150 mM NaCl). GPCRs are known to bind anionic headgroup containing phospholipids and this binding interaction is believed to have roles in GPCR activation and G-protein binding (29–31). Competitive membrane binding interactions could significantly influence the binding of TIL to its protein binding partners *in vivo*. Therefore, we asked if the TIL-SH3 interaction would be influenced by the interactions of TIL and anionic lipid headgroups. For this experiment, we used endophilin-SH3 domain since the full length endophilin would bind to the negatively charged membranes irrespective of TIL-SH3 interactions. The purified SH3 domain was labeled at the N-terminus with Alexa 594-NHS ester for fluorescence microscopy imaging. Interestingly, the SH3 domain did not show detectable binding to the GUV surface with electrostatically bound TIL. In contrast, TIL conjugated to neutral membranes allowed recruitment of SH3 (Figure 3C, 3D). Our data therefore indicate that the TIL-membrane interaction could be a crucial regulator of SH3 domain mediated endophilin recruitment via TIL.

**Figure 3.**
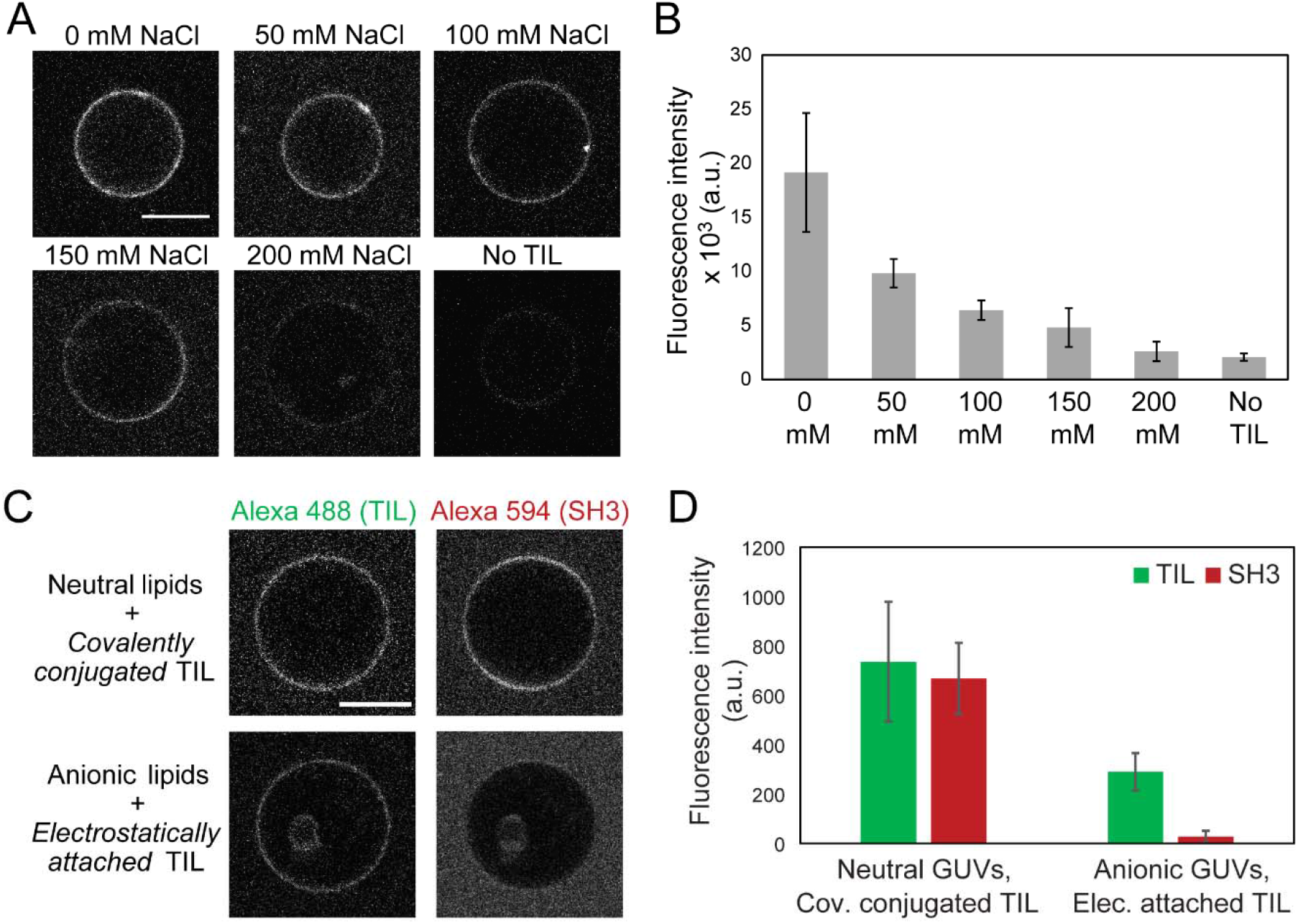
TIL spontaneously binds to anionic lipid containing membranes, and the TIL-anionic lipid interaction interferes with SH3-domain recruitment. A. Confocal images showing TIL-Alexa 488 fluorescence from the GUVs composed of DOPS/DOPE/DOPC (45:30:25). The ionic strength of the bulk solution was varied by changing the sodium chloride content in buffer (20 mM HEPES, pH 7.4) from 0-200 mM. B. Bar plot showing the fluorescence intensity of bound TIL-Alexa 488 on GUVs at various salt concentrations. A decrease in the binding with increasing ionic strength suggests that the nature of binding is electrostatic. C. Confocal images showing recruitment of TIL and SH3 on GUVs when TIL is covalently conjugated to GUVs composed of neutral lipids (mCC-PE/PC 5:95) vs. when TIL is electrostatically attached to GUVs containing anionic lipids (PS/PE/PC 45:30:25). D. Bar plot showing the fluorescence intensities in the Alexa 488 (TIL) channel and Alexa 594 (SH3) channel from the GUV surfaces indicates that covalently coupled TIL recruits the SH3 domain whereas electrostatically attached TIL does not. All scale bars are 10 µm. Bar plots are represented as mean ± s.d, N ≥ 6 for 3B and N ≥ 11 for 3D.

### Endophilin tubulates neutral membranes when recruited by TIL

Endophilin has been widely investigated for its capacity to tubulate lipid bilayers in vitro and when overexpressed in live cells. Both full endophilin and the N-BAR domains are known to tubulate LUVs composed of anionic phospholipids, as demonstrated earlier with transmission electron microscopy (TEM) (18, 19). Mechanisms of curvature generation by BAR-domain proteins that have been suggested include 1) scaffolding – generation of a 3D protein assembly upon oligomerization of the membrane-bound BAR-proteins that modulate membrane shape (32), 2) amphipathic helix insertion – embedding of the N-terminal amphipathic helix present in the N-BAR sub-family proteins resulting in membrane bending (20), and 3) protein crowding – enhancement of membrane spontaneous curvature in order to reduce the steric pressure generated by crowded membrane anchored proteins (33, 34). These mechanisms are based on observations that these proteins primarily anchor to the membrane via the N-BAR domain that forms strong electrostatic interactions with the anionic lipid headgroups. We therefore asked whether membrane curvature could be induced by recruitment of endophilin to the membrane primarily through the SH3 domain-mediated interaction. In order to answer this question, we prepared LUVs of 400 nm diameter (*Supplementary Information*, Figure S4) containing maleimide lipids, to which we covalently coupled TIL, followed by incubation with endophilin. Interestingly, TEM images show that 35% of the TIL conjugated LUVs form tubules in the presence of endophilin. Only about 2% of the TIL functionalized LUVs show tubulation in the absence of endophilin. LUVs without TIL conjugation do not show any tubulation in the presence of endophilin (Figure 4A, 4B).

**Figure 4.**
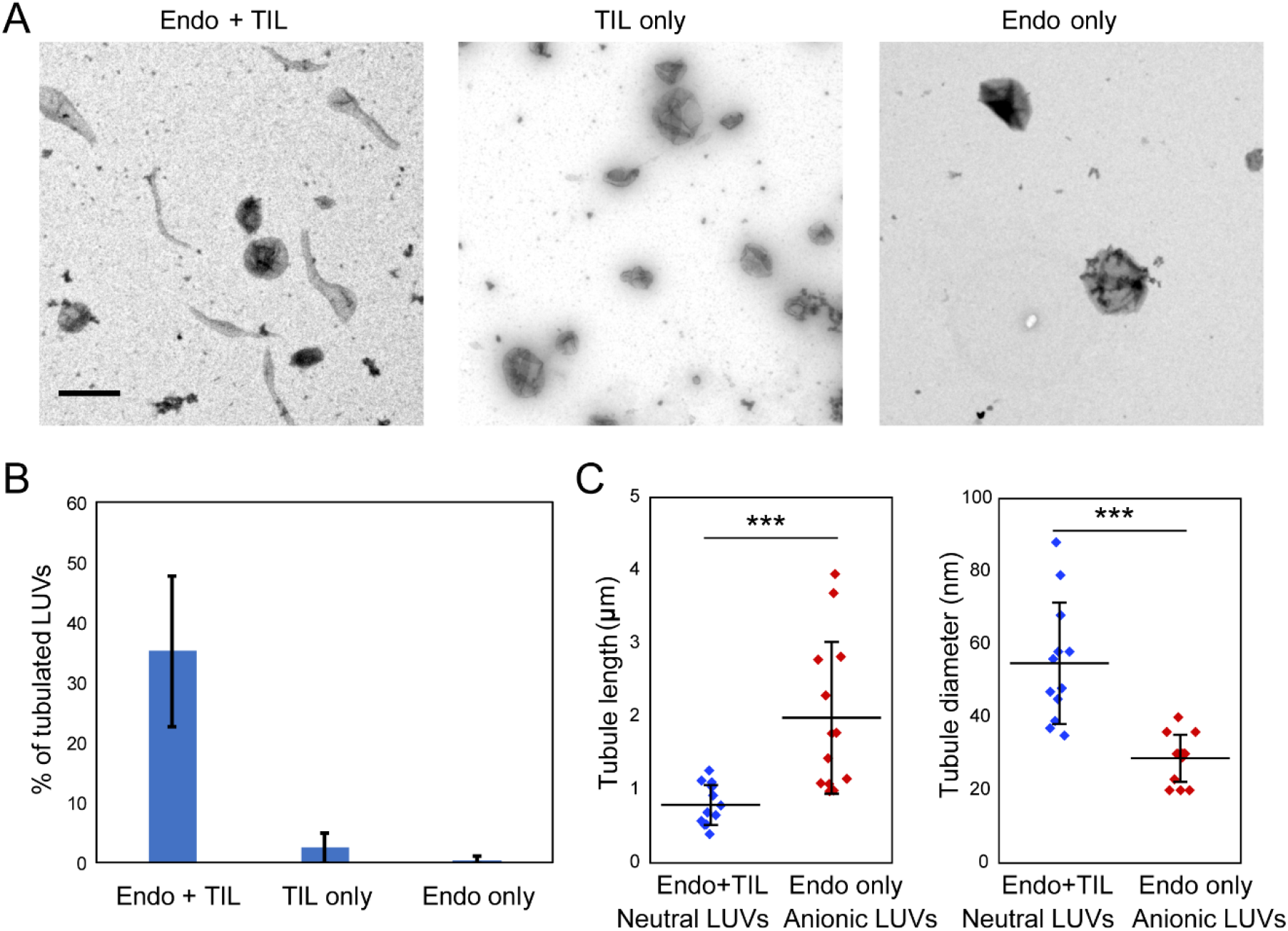
Endophilin tubulates TIL-conjugated LUVs even in the absence of anionic phospholipids. A. TEM images of LUVs composed of 5% MCC-PE and 95% DOPC conjugated with TIL shows tubulation in the presence of endophilin (left panel) whereas there is no tubulation in case of TIL-conjugated LUVs in the absence of endophilin (middle panel). Conversely, endophilin itself does not tubulate LUVs that are not conjugated with TIL (right panel). Scale bar 500 nm. B. Bar plot showing the percentage of tubulated, vesicles in the presence and absence of endophilin. Error bar ± s.d. C. Scatter plots showing the distribution of tubule length and diameter formed by endophilin with TIL-conjugated LUVs (blue) and with LUVs composed of anionic phospholipids (DOPS:DOPE:DOPC 45:30:25) (red). The horizontal lines indicate mean values of the distributions and the whiskers indicate standard deviations. Student’s t-test, *** *P* ≤ 0.001, *N* = 13.

We determined the tubule diameters from the TEM images in the case of TIL-recruited endophilin and compared them with the tubules obtained in case of anionic lipid recruited endophilin (Figure 4C, and *Supplementary Information*, Figure S5). Identical endophilin:lipid ratios and incubation times were maintained for both cases. Tubule diameters on anionic lipid containing membranes were in the range of 20-40 nm, comparable with earlier reports in the presence of endophilin N-BAR domain (20). The tubule diameters were significantly larger, in the range of 40-80 nm, in the case of TIL-recruited endophilin to membranes having no anionic lipids (Figure 4C). Furthermore, the tubules generated by TIL-recruited endophilin were about 1 µm or shorter in length whereas the length of the tubules formed in the case of endophilin with anionic lipids were in the range of 1-5 µm. These observations imply that the curvature generation is higher for the self-recruited endophilin onto negatively membranes. Our results imply that endophilin can deform membranes when recruited to membranes exclusively by TIL.

### TIL-recruited endophilin is curvature sorted on membranes with neutral net-charge

BAR domain proteins sense membrane curvature through preferential partitioning into highly bent membrane regions (22, 35–37). Endophilin has been shown to be recruited at the sites of membrane invagination during CIE and promote membrane scission (15). Our previous studies have shown that endophilin N-BAR is sorted onto membrane tethers pulled from GUVs composed of anionic phospholipids (19, 22). Next, we asked if this curvature sorting behavior of endophilin is observed when the protein is bound to membranes with neutral net-charge through membrane-bound TIL. Incorporation of a small (0.2 mol%) amount of lipids containing a biotin-functionalized headgroup facilitated the pulling of membrane tethers from GUVs with the help of streptavidin-functionalized beads (22). When tethers were pulled from TIL-conjugated GUVs in the presence of endophilin, we observed significantly higher fluorescence intensities of endophilin on tethers compared to the flatter membrane surface indicating curvature sorting of endophilin (Figure 5A, 5B and 5E).

**Figure 5.**
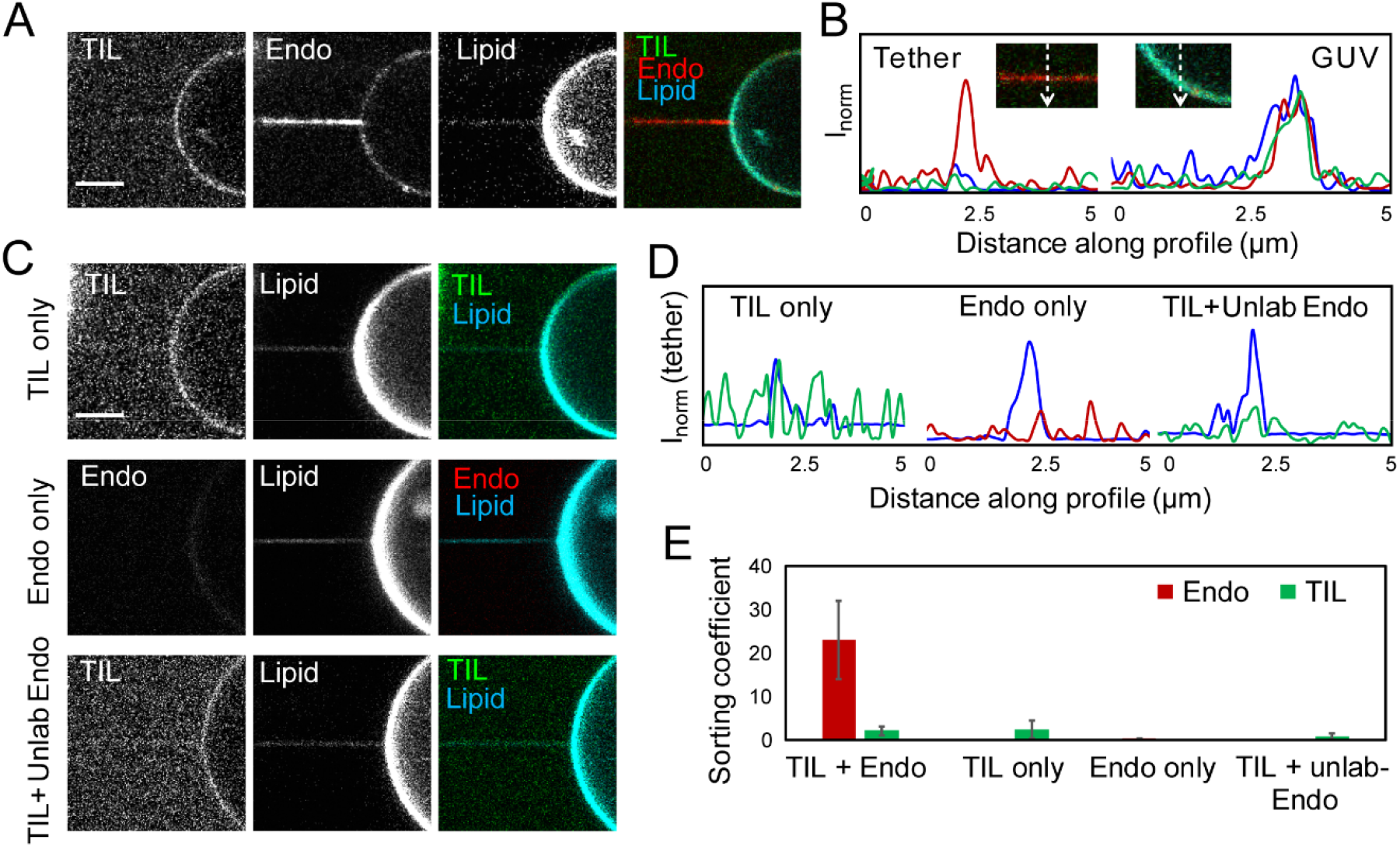
Curvature sorting of TIL-recruited endophilin observed in membranes with neutral net-charge. A. Confocal fluorescence images showing a membrane tether pulled from a GUV conjugated to TIL-Alexa 488 in the presence of endophilin-Alexa 594. Images are recorded in three different channels to show the distribution of TIL, endophilin, and the lipid dye DiD on the membrane tether and on the flatter GUV area. In the endophilin channel, the tether shows higher fluorescence intensity compared to the vesicle, indicating curvature sorting of the protein. Scale bar 5 μm. B. Intensity profiles of endophilin (red), TIL (green), and DiD (blue) on the membrane tether and on the GUV surface along the white lines shown in the inset. Higher intensity in the red channel from the tether indicates endophilin is enriched in the membrane tether region compared to the flatter GUV area. C. Confocal images showing tether pulled from a TIL-Alexa 488 conjugated GUV in the absence of endophilin (TIL only), top panel; from a GUV having no TIL-Alexa 488 conjugated in the presence of endophilin (Endo only), middle panel; and from a TIL-Alexa 488 conjugated GUV in the presence of unlabeled endophilin (TIL+unlab Endo), bottom panel. Scale bar 5 µm. D. Intensity profiles across the tether region of the images shown in Figure 5C. E. Bar plot showing sorting coefficients for endophilin and TIL as quantified from the corresponding fluorescence images under the conditions mentioned in Figure 5B and 5C using the formula described in the methods section. Error bars represent standard error of means calculated from 4 independent trials, at least 3 GUVs were studied in each trial.

Notably, no such curvature sorting was observed in the TIL channel (Figure 5A, 5B, and 5E). One possibility behind not observing curvature sorting in the TIL channel could be loss of Alexa 488 fluorescence signal due to energy transfer to the Alexa 594-labels on neighboring endophilin molecules. To exclude this possibility, we pulled tethers from TIL-conjugated GUVs in the presence of unlabeled endophilin (Figure 5C bottom panel). No curvature sorting of TIL was observed in this case either (Figure 5D, 5E). These data indicate that TIL-bound endophilin, but not TIL on its own, is curvature sorted. Furthermore, tethers pulled from TIL-conjugated GUVs in the absence of endophilin also did not show any curvature sorting of TIL indicating that the accumulation of endophilin on tethers is not due to curvature sorting of TIL (Figure 5C top panel, 5D and 5E). A second hypothesis is that the conjugated TIL is immobile on the membrane. We tested this second hypothesis through fluorescence recovery after photobleaching experiments that demonstrated recovery of TIL-Alexa 488 fluorescence from 60% to 90% in 20 s (*Supplementary Information*, Figure S6). Therefore, TIL clearly is mobile on the membrane. Given these observations, we hypothesize that under our experimental conditions, TIL is too crowded on the membrane to show significant curvature sorting (22). Finally, in the absence of TIL, no curvature sorting of endophilin was observed since by itself, it does not show binding to the uncharged membrane (Figure 5C middle panel, 5D and 5E).

Taken together, these findings strongly suggest that the curvature sensing capabilities of endophilin persist during its recruitment to the membrane by the SH3-TIL interaction even if the membrane does not contain anionic phospholipids.

## Discussion

Electrostatic interactions between BAR-domains and anionic lipid headgroups are considered critical for curvature sensing and generation by BAR-proteins (20, 38–41). While hydrophobic interactions have also been shown to play significant roles in curvature generation, electrostatic contributions have not been separated in those studies (39). Therefore, to what extent electrostatic interactions contribute to the curvature sensing and generation by BAR domains has remained unclear. A major challenge faced is the loss of membrane binding of BAR-domain proteins in the absence of long-range electrostatic attractions provided by anionic lipids and the cationic residues of the BAR domain. To overcome this challenge, we covalently conjugated the TIL of a GPCR to the membrane. Our observation that the BAR domain retains its curvature sensing and (some of its) curvature generation properties in the absence of long-range electrostatic interactions is striking. Short range, non-specific interactions such as hydrophobic interactions, cation-π interactions, and H-bonding interactions with zwitterionic choline headgroups have previously been shown to be important in the recruitment of various peripheral proteins to uncharged membranes (42–45). How they are involved in curvature sensing and generation by BAR domains is important to understand.

### Proposed mechanism of curvature sensing and generation on membranes with neutral-net charge

Models based on results from electron microscopy imaging, electron paramagnetic resonance spectroscopy, and molecular dynamics simulations suggested that BAR-domains adopt a specific orientation with respect to the membrane while sensing and generating membrane curvature (18, 21, 36). In this orientation, the concave surface of the BAR domain faces toward the membrane surface stabilized by electrostatic interactions between cationic residues clustered at the concave surface and anionic lipid headgroups. This orientation also allows membrane insertion of hydrophobic residues present at the interior of the concave surface that have been found to be crucial for curvature generation (39). We hypothesize that in addition to electrostatic interactions, hydrophobic effects, along with cation-π interactions and H-bonding interactions, play crucial roles in orienting the BAR domain with respect to the membrane as well. Apparently, the combined effect of these interactions is strong enough to drive curvature sensing and generation even in the absence of long-range electrostatic interactions.

In peripheral proteins, hydrophobic residues are considered to reinforce membrane bound states by penetrating into the hydrocarbon core unlike charged residues that merely interact with the lipid headgroups (42, 44). Amphipathic peptides having cationic and hydrophobic sidechains have been shown to sense membrane curvature by their insertion into the membrane through lipid packing defects (36, 46). Highly curved membranes are considered to contain more lipid packing defects and hence are thought to facilitate hydrophobic insertion. In the case of endophilin N-BAR domains, insertion of its two amphipathic helix forming regions has been considered to be crucial for curvature sensing and generation, but their actual roles are highly debated (18, 20, 36, 47). Cryo EM and simulation studies have suggested involvement of amphipathic helices in protein-scaffold formation on tubular membranes (21, 48). On the other hand, a previous contribution from our group (consistent with results from Fernandes et al. (49)) showed that insertion of the N-terminal amphipathic region is not necessary for either curvature sensing or curvature generation by the endophilin BAR domain, although this region is crucial for membrane binding (19). These findings suggest that rather than the H0 helix dominating endophilin’s function in sensing and generating membrane curvature, the crescent shaped BAR domain appears to play a critical role in these phenomena.

Among hydrophobic residues present on the BAR domain surface, Leu166 has been found by EPR studies to be inserted into the bilayer beyond the phosphate level of membrane tubules (18). From the crystal structure of the dimeric N-BAR domain (PDB: 2C08), we evaluated the surface distributions of such non-polar amino acid sidechains which, upon insertion into the membrane, might help the BAR domain adopt a specific orientation with respect to the membrane. Non-polar aliphatic residues Met, Cys, Leu, and Ile along with the aromatic sidechains were shown by Wimley and White to offer a favorable free energy of partitioning from the aqueous phase to the membrane interface (50). Both the concave and the convex surfaces of the dimeric BAR domain contain six hydrophobic sidechains along with two aromatic sidechains that have more than 50% surface accessibility as determined by the GETAREA tool (51) (Figure 6A). Neither the concave nor the convex surface would allow all six hydrophobic residues to interact simultaneously with a flat membrane surface. (Figure 6B and *Supplementary Information*, Figures S7 A, B). However, on a membrane with positive curvature, the concave surface would allow all six hydrophobic residues to interact with the membrane whereas the convex surface would allow a maximum of three hydrophobic interactions (Figure 6B and *Supplementary Information*, Figure S7 C). This indicates that hydrophobic interactions would be most favorable on a curved membrane if the protein binds through its concave surface.

**Figure 6.**
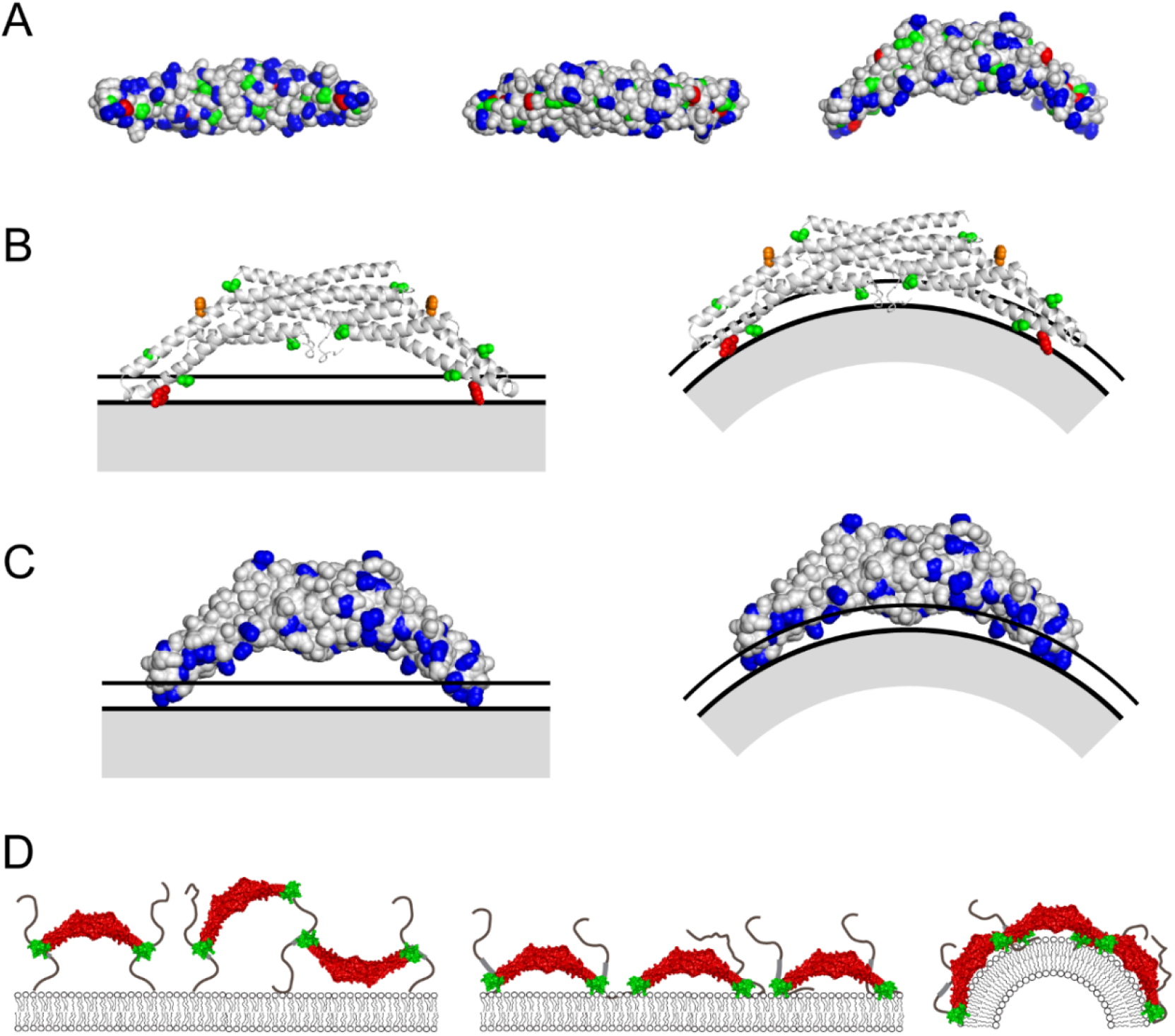
Short range interactions facilitate curvature sensing and generation by BAR domain during TIL-mediated recruitment. A. Solvent accessible surface of endophilin BAR domain (PDB: 2C08) viewed from the bottom (left), top (middle) and side (right). Sidechains of cationic, hydrophobic aliphatic, and aromatic residues are highlighted in blue, green, and red respectively. B. Interaction of the surface accessible aromatic and aliphatic residues with non-polar sidechains with flat (left) and curved (right) membranes. Solid line: choline headgroups, dotted line: phosphate region, grey area: lipid tail region. The BAR domain is shown in cartoon representation so that the hydrophobic residues of interest are clearly visible. Aliphatic hydrophobic residues with 50% or more solvent accessible surface area are shown as green spheres. Aromatic residues that can form additional cation-π interactions with choline headgroups are shown in different colors (red spheres: Tyr, orange spheres: Phe). C. Interaction of the accessible cationic sidechains present at the concave surface of BAR-domain with the lipid phosphate groups with flat (left) and curved (right) membranes. D. Mechanism of curvature generation and sensing by TIL-recruited full length endophilin (N-BAR domain in red and SH3 domain in green). The SH3 domains are placed at the distal tips of the N-BAR dimer following the structural model described in ref. (60)). TIL-SH3 interactions help initial anchoring of endophilin to the membrane surface (left) followed by the short-range interactions that help re-orienting the BAR-domain with respect to the membrane (middle). Suitably oriented BAR domains facilitate membrane curvature generation and larger extent of stabilizing interactions on curved membranes drive curvature sorting of endophilin (right).

Among the aromatic residues, Tyr in particular has been shown to form cation-π interactions with zwitterionic PC headgroups that can provide a stabilizing free energy contribution of up to 3 kcal/mol (45, 52). The endophilin BAR domain has two Tyr170 near the two tips of the dimeric BAR domain on the concave surface. Both of these tyrosines would stabilize the concave surface facing the membrane by interacting with the choline headgroups. This orientation would favor the cation-π interactions on both flat and curved membrane surfaces (Figure 6B).

Cationic residues in proteins are found to form H-bonding interactions with the phosphates of lipid headgroups (43, 53). How they contribute to the protein binding with zwitterionic PC lipids is under debate. While Wimley and White, using synthetic pentapeptides, have shown that the experimentally determined ΔG is unfavorable to the transfer of Lys and Arg from solution to the membrane interface, computational work by MacCallum et al. suggests a negative ΔG at the membrane interface for small molecule analogs of cationic Lys and Arg sidechains (50, 54). Additionally, experimental studies with poly-lysine and poly-arginine peptides have not only found binding with purely zwitterionic membranes but also observed membrane deformation by the cationic residues (53, 55). The modes of interaction are believed to be electrostatic, and to also include H-bonding, with possible contributions from hydrophobic interactions as well (53, 56). Also, it has been shown in the case of peripheral proteins such as phosphatidylinositol-specific phospholipase C, that the removal of a key H-bond forming lysine residue reduces affinity to PC membranes (43). A majority of the cationic residues are present on the concave surface of the BAR domain (Figure 6A). Therefore, orientation of this surface toward the membrane would allow a maximal number of the cationic sidechains to interact with the PC lipids compared to the other orientations (Figure 6C and *Supplementary Information*, Figure S7 A). Adopting this orientation on curved membranes would bring most of the cationic sidechains from the concave surface closer to the lipid headgroups. These interactions, besides helping in curvature sensing, could also contribute to membrane deformation by the BAR domain in a similar manner as polycationic peptides.

Our study shows that the short-range interactions, in a nanomolar concentration range of the BAR domain, are incapable of effectively recruiting endophilin to the uncharged membrane in the absence of TIL. However, when the protein is recruited to the membrane via TIL-SH3 interaction, the local concentration of endophilin on the membrane increases. Under such conditions, the short-range interactions would become significant for orienting the protein such that the cationic residue rich concave surface faces the membrane. We hypothesize that the initial anchoring via the flexible TIL would allow various orientations of the protein on the membrane surface (Figure 6D). The effect of short-range interactions would then lead to an orientation where the concave surface faces the membrane.

Short range interactions with the concave surface are maximized on curved membranes. Therefore, the membrane residence time of the BAR domain would be longer for a cylindrically shaped membrane over a flat membrane. This hypothesis is supported by our observation that TIL recruited endophilin is curvature sorted onto membrane tethers. Furthermore, we showed that in the absence of anionic lipids, endophilin causes tubulation of LUVs although to a lesser extent compared to anionic membranes. Overall, our study shows that while long range electrostatic interactions could be important for the recruitment of endophilin to the membrane, short range interactions are crucial to drive curvature sensing and generation.

Whereas endophilin can sense and generate curvature in the presence of anionic lipids in vitro, the membrane remodeling activity of endogenous endophilin in cells in the FEME pathway requires activation of GPCR followed by heterotrimeric G-protein release (10, 14), which makes available for endophilin binding the GPCR’s TIL. We have shown that TIL binding not only recruits endophilin to charge-neutral membranes but also facilitates membrane bending, a key step in receptor internalization. A key aspect revealed here is that endophilin’s membrane remodeling activity is not limited to negatively charged membranes. These observations raise the question whether the electrostatic interactions between positively charged residues on the N-BAR domain and negatively charged lipid headgroups primarily mediate protein recruitment to the membrane or if they play a critical role in membrane bending as well.

Our finding that anionic lipid headgroups interfere with TIL binding to SH3-domains indicates that recognition of the activated GPCR by endophilin could be regulated by the lipid microenvironment of the receptor. We speculate that molecular interactions between BAR-proteins, GPCR-TIL, and the phospholipid headgroups play an essential role in the downregulation of GPCRs by endocytic pathways.

## Materials and Methods

A brief description of experimental methods is given below. Detailed Materials and Methods with additional figures are provided in the *Supplementary Information*.

The TIL of human β1-AR was cloned into an E. coli expression vector to allow expression as an N-terminally GST tagged protein. After overexpression, the GST-tagged TIL was purified from E. coli cell lysate using GST-affinity chromatography. The GST-tag was cleaved by protease treatment and TIL was further purified by cation exchange chromatography. N-terminal labeling of TIL was performed using Alexa 488-SDP ester in 10 mM phosphate buffer at pH 7.0 and the unreacted fluorophores were removed by desalting columns.

The single cysteine mutants of full length rat endophilin A1 and its N-BAR domain were purified and labeled with Alexa 594 maleimide-derivatives following earlier reported procedures (57). The SH3 domain was also purified following the same protocol as full length protein and was labeled at the N-terminus with Alexa 594-NHS ester.

GUVs were prepared by the electroformation method (58). TIL-conjugation was performed by incubating the GUVs with TIL in HEPES buffer containing NaCl and TCEP, pH 7.4 at room temperature for 6-8 hours. LUVs were prepared by the extrusion and TIL conjugation of LUVs were performed following the same procedure as used for GUVs.

All confocal images were recorded using lass coverslips passivated with β-casein. Fluorescence signals from Alexa 488, Alexa 594, and DiD were recorded at three independent detectors upon sequential excitation of the fluorophores using three different lasers emitting at 488 nm, 560 nm, and 640 nm respectively.

For membrane tether pulling, GUVs containing biotin-functionalized lipids were aspirated with a glass capillary passivated with β-casein. Another capillary was used to capture a streptavidin coated bead (6 µm diameter). Both the capillaries were controlled by the motorized arms of a micromanipulator instrument. The bead was touched to the GUV surface and then moved far from the surface to form a thin membrane tether. Sorting coefficients were calculated from the intensities determined at the protein and lipid channels for tether and flat GUV regions using the formula (59):

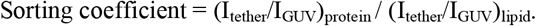

LUV tubulation assays were performed by incubating the LUVs with endophilin in HEPES buffer for 30 minutes at room temperature. To prepare samples for negative staining TEM imaging, LUV samples adhered on carbon coated copper grids were stained using 2% uranyl acetate solution and excess stains were removed by filter papers. Grids were further washed with buffer and air-dried.

## Acknowledgments

The authors thank Jaclyn Robustelli and Rui Jin for generously providing purified proteins. We are grateful to Tatyana Svitkina lab for the TEM facility. We acknowledge helpful comments and suggestions from Zachary Zimmerman and Sankalp Shukla. This work was financially supported by National Institute of Health, grant No GM 097552.

## Author Contributions

S.M. and T.B. designed research. S.M. performed research. S.M., I.P., K.N., and S.B. contributed in data analysis. S.M. and T.B. wrote the paper.

## Notes

The authors declare no competing interest.

## References

1. S. S. Ferguson, Evolving concepts in G protein-coupled receptor endocytosis: the role in receptor desensitization and signaling. Pharmacol Rev 53, 1–24 (2001).

2. R. Irannejad, M. von Zastrow, GPCR signaling along the endocytic pathway. Curr Opin Cell Biol 27, 109–116 (2014).

3. A. Sorkin, M. Von Zastrow, Signal transduction and endocytosis: close encounters of many kinds. Nature reviews. Molecular cell biology 3, 600–614 (2002).

4. V. Vieira, C. Lamaze, S. L. Schmid, Control of EGF receptor signaling by clathrin-mediated endocytosis. Science 274, 2086–2089 (1996).

5. S. D. Conner, S. L. Schmid, Regulated portals of entry into the cell. Nature 422, 37–44 (2003).

6. M. Kaksonen, A. Roux, Mechanisms of clathrin-mediated endocytosis. Nature reviews. Molecular cell biology 19, 313–326 (2018).

7. H. T. McMahon, E. Boucrot, Molecular mechanism and physiological functions of clathrin-mediated endocytosis. Nature reviews. Molecular cell biology 12, 517–533 (2011).

8. Y. Mosesson, G. B. Mills, Y. Yarden, Derailed endocytosis: an emerging feature of cancer. Nature reviews. Cancer 8, 835–850 (2008).

9. A. Frost, V. M. Unger, P. De Camilli, The BAR domain superfamily: membrane-molding macromolecules. Cell 137, 191–196 (2009).

10. P. A. Ferreira, E. Boucrot, Mechanisms of Carrier Formation during Clathrin-Independent Endocytosis. Trends Cell Biol 28, 188–200 (2018).

11. G. Cestra et al., The SH3 domains of endophilin and amphiphysin bind to the proline-rich region of synaptojanin 1 at distinct sites that display an unconventional binding specificity. The Journal of biological chemistry 274, 32001–32007 (1999).

12. N. Ringstad, Y. Nemoto, P. De Camilli, Differential expression of endophilin 1 and 2 dimers at central nervous system synapses. The Journal of biological chemistry 276, 40424–40430 (2001).

13. L. Bertot et al., Quantitative and Statistical Study of the Dynamics of Clathrin-Dependent and -Independent Endocytosis Reveal a Differential Role of EndophilinA2. Cell Rep 22, 1574–1588 (2018).

14. E. Boucrot et al., Endophilin marks and controls a clathrin-independent endocytic pathway. Nature 517, 460–465 (2015).

15. H. F. Renard et al., Endophilin-A2 functions in membrane scission in clathrin-independent endocytosis. Nature 517, 493–496 (2015).

16. Y. Tang et al., Identification of the endophilins (SH3p4/p8/p13) as novel binding partners for the beta1-adrenergic receptor. Proc Natl Acad Sci U S A 96, 12559–12564 (1999).

17. A. Casamento, E. Boucrot, Molecular mechanism of Fast Endophilin-Mediated Endocytosis. Biochemical Journal 477, 2327–2345 (2020).

18. M. R. Ambroso, B. G. Hegde, R. Langen, Endophilin A1 induces different membrane shapes using a conformational switch that is regulated by phosphorylation. Proc Natl Acad Sci U S A 111, 6982–6987 (2014).

19. Z. Chen, C. Zhu, C. J. Kuo, J. Robustelli, T. Baumgart, The N-Terminal Amphipathic Helix of Endophilin Does Not Contribute to Its Molecular Curvature Generation Capacity. J Am Chem Soc 138, 14616–14622 (2016).

20. J. L. Gallop et al., Mechanism of endophilin N-BAR domain-mediated membrane curvature. The EMBO journal 25, 2898–2910 (2006).

21. C. Mim et al., Structural basis of membrane bending by the N-BAR protein endophilin. Cell 149, 137–145 (2012).

22. C. Zhu, S. L. Das, T. Baumgart, Nonlinear sorting, curvature generation, and crowding of endophilin N-BAR on tubular membranes. Biophys J 102, 1837–1845 (2012).

23. K. Farsad et al., Generation of high curvature membranes mediated by direct endophilin bilayer interactions. The Journal of cell biology 155, 193–200 (2001).

24. R. J. Brea et al., In Situ Reconstitution of the Adenosine A2A Receptor in Spontaneously Formed Synthetic Liposomes. J Am Chem Soc 139, 3607–3610 (2017).

25. T. Kimura et al., Recombinant cannabinoid type 2 receptor in liposome model activates g protein in response to anionic lipid constituents. The Journal of biological chemistry 287, 4076–4087 (2012).

26. L. Niu, J. M. Kim, H. G. Khorana, Structure and function in rhodopsin: asymmetric reconstitution of rhodopsin in liposomes. Proc Natl Acad Sci U S A 99, 13409–13412 (2002).

27. E. Serebryany, G. A. Zhu, E. C. Yan, Artificial membrane-like environments for in vitro studies of purified G-protein coupled receptors. Biochimica et biophysica acta 1818, 225–233 (2012).

28. M. Brinkley, A brief survey of methods for preparing protein conjugates with dyes, haptens and crosslinking reagents. Bioconjugate chemistry 3, 2–13 (1992).

29. R. Dawaliby et al., Allosteric regulation of G protein-coupled receptor activity by phospholipids. Nat Chem Biol 12, 35–39 (2016).

30. W. Song, H. Y. Yen, C. V. Robinson, M. S. P. Sansom, State-dependent Lipid Interactions with the A2a Receptor Revealed by MD Simulations Using In Vivo-Mimetic Membranes. Structure 27, 392–403 e393 (2019).

31. H. Y. Yen et al., PtdIns(4,5)P2 stabilizes active states of GPCRs and enhances selectivity of G-protein coupling. Nature 559, 423–427 (2018).

32. M. Simunovic et al., How curvature-generating proteins build scaffolds on membrane nanotubes. Proc Natl Acad Sci U S A 113, 11226–11231 (2016).

33. W. T. Snead et al., Membrane fission by protein crowding. Proc Natl Acad Sci U S A 114, E3258–E3267 (2017).

34. W. T. Snead, J. C. Stachowiak, Structure Versus Stochasticity-The Role of Molecular Crowding and Intrinsic Disorder in Membrane Fission. J Mol Biol 430, 2293–2308 (2018).

35. B. Antonny, “Mechanisms of Membrane Curvature Sensing” in Annual Review of Biochemistry, Vol 80, R. D. Kornberg, C. R. H. Raetz, J. E. Rothman, J. W. Thorner, Eds. (2011), vol. 80, pp. 101–123.

36. H. Cui, E. Lyman, G. A. Voth, Mechanism of membrane curvature sensing by amphipathic helix containing proteins. Biophys J 100, 1271–1279 (2011).

37. H. T. McMahon, E. Boucrot, Membrane curvature at a glance. J Cell Sci 128, 1065–1070 (2015).

38. J. Peter et al., BAR domains as sensors of membrane curvature: the amphiphysin BAR structure. Science 303, 495–499 (2004).

39. M. Masuda et al., Endophilin BAR domain drives membrane curvature by two newly identified structure-based mechanisms. The EMBO journal 25, 2889–2897 (2006).

40. Y. Yoon, X. Zhang, W. Cho, Phosphatidylinositol 4,5-bisphosphate (PtdIns(4,5)P2) specifically induces membrane penetration and deformation by Bin/amphiphysin/Rvs (BAR) domains. The Journal of biological chemistry 287, 34078–34090 (2012).

41. J. Zimmerberg, S. McLaughlin, Membrane curvature: how BAR domains bend bilayers. Current biology : CB 14, R250–252 (2004).

42. N. Ben-Tal, B. Honig, C. Miller, S. McLaughlin, Electrostatic binding of proteins to membranes. Theoretical predictions and experimental results with charybdotoxin and phospholipid vesicles. Biophys J 73, 1717–1727 (1997).

43. H. M. Khan et al., A Role for Weak Electrostatic Interactions in Peripheral Membrane Protein Binding. Biophys J 110, 1367–1378 (2016).

44. A. L. Lomize, I. D. Pogozheva, M. A. Lomize, H. I. Mosberg, The role of hydrophobic interactions in positioning of peripheral proteins in membranes. BMC Struct Biol 7, 44 (2007).

45. Q. Waheed et al., Interfacial Aromatics Mediating Cation-pi Interactions with Choline-Containing Lipids Can Contribute as Much to Peripheral Protein Affinity for Membranes as Aromatics Inserted below the Phosphates. The journal of physical chemistry letters 10, 3972–3977 (2019).

46. N. S. Hatzakis et al., How curved membranes recruit amphipathic helices and protein anchoring motifs. Nat Chem Biol 5, 835–841 (2009).

47. V. K. Bhatia et al., Amphipathic motifs in BAR domains are essential for membrane curvature sensing. The EMBO journal 28, 3303–3314 (2009).

48. H. Cui et al., Understanding the role of amphipathic helices in N-BAR domain driven membrane remodeling. Biophys J 104, 404–411 (2013).

49. F. Fernandes et al., Role of helix 0 of the N-BAR domain in membrane curvature generation. Biophys J 94, 3065–3073 (2008).

50. W. C. Wimley, S. H. White, Experimentally determined hydrophobicity scale for proteins at membrane interfaces. Nat Struct Biol 3, 842–848 (1996).

51. R. Fraczkiewicz, W. Braun, Exact and efficient analytical calculation of the accessible surface areas and their gradients for macromolecules. Journal of computational chemistry 19, 319–333 (1998).

52. Y. Hirano et al., Structural basis of phosphatidylcholine recognition by the C2-domain of cytosolic phospholipase A2alpha. Elife 8(2019).

53. A. D. Robison et al., Polyarginine Interacts More Strongly and Cooperatively than Polylysine with Phospholipid Bilayers. The journal of physical chemistry. B 120, 9287–9296 (2016).

54. J. L. MacCallum, W. F. Bennett, D. P. Tieleman, Distribution of amino acids in a lipid bilayer from computer simulations. Biophys J 94, 3393–3404 (2008).

55. T. A. Spurlin, A. A. Gewirth, Poly-L-lysine-induced morphology changes in mixed anionic/zwitterionic and neat zwitterionic-supported phospholipid bilayers. Biophys J 91, 2919–2927 (2006).

56. M. Hoernke, C. Schwieger, A. Kerth, A. Blume, Binding of cationic pentapeptides with modified side chain lengths to negatively charged lipid membranes: Complex interplay of electrostatic and hydrophobic interactions. Biochimica Et Biophysica Acta (BBA)-Biomembranes 1818, 1663–1672 (2012).

57. B. R. Capraro et al., Kinetics of endophilin N-BAR domain dimerization and membrane interactions. The Journal of biological chemistry 288, 12533–12543 (2013).

58. L. Mathivet, S. Cribier, P. F. Devaux, Shape change and physical properties of giant phospholipid vesicles prepared in the presence of an AC electric field. Biophys J 70, 1112–1121 (1996).

59. B. Sorre et al., Curvature-driven lipid sorting needs proximity to a demixing point and is aided by proteins. Proc Natl Acad Sci U S A 106, 5622–5626 (2009).

60. Q. Wang, H. Y. K. Kaan, R. N. Hooda, S. L. Goh, H. Sondermann, Structure and Plasticity of Endophilin and Sorting Nexin 9. Structure 16, 1574–1587 (2008).

