## Supplementary Information for "Endophilin Recruitment via GPCR Interactions Enables Membrane Curvature Generation in the Absence of Anionic Lipids"

### Materials

The lipids 1,2-dioleoyl-sn-glycero-3-phosphocholine (DOPC), 1,2-dioleoyl-sn-glycero-3-phosphoethanolamine (DOPE), 1,2-dioleoyl-sn-glycero-3-phospho-L-serine (DOPS), 1,2-dipalmitoyl-sn-glycero-3-phosphoethanolamine-N-[4-(p-maleimidomethyl)cyclohexane-carboxamide] (16:0 mCC-PE), and 1,2-distearoyl-sn-glycero-3-phosphoethanolamine-N-[biotinyl(polyethylene glycol)-2000] (DSPE-PEG-Biotin) were obtained from Avanti Polar Lipids (Alabaster, AL). Alexa Fluor 488 sulfodichlorophenol (SDP) ester, Alexa Fluor 594 C5 maleimide, and the lipid fluorophore 1,1'-dioctadecyl-3,3,3',3'-tetramethylindodicarbocyanine, 4-chlorobenzenesulfonate salt (DiD) were from ThermoFisher Scientific (USA). Streptavidin conjugated microsphere beads (6 µm diameter) were from Polysciences (Warrington, PA). All common reagents used for the buffer preparation, such as HEPES, Tris, NaCl, $Na_2HPO_4$, $NaH_2PO_4$, ethylenediaminetetraacetic acid (EDTA), and dithiothreitol (DTT) were from Fisher Scientific (USA). β-casein from bovine milk was obtained from Sigma-Aldrich (USA). Unless otherwise specified, all chemicals from commercial sources were used without further purification.

### Methods

**Construct design, protein expression, and purification of TIL.** The TIL region of the β1-adrenergic receptor (amino acids 246-325) was cloned from a pcDNA3-Flag beta-1-adrenergic-receptor vector (Robert Lefkowitz lab, Addgene Plasmid #14698) by PCR using the following primers: forward 5'-AAAGGATCCCGGGTGTTCCGCG-3' and reverse 5'-AAAGAATTCTTACGTCTTGAGCGCCTTCTG-3'). The TIL sequence was ligated into BamHI/EcoRI restriction sites of the pGEX6p1 vector for bacterial expression as an N-terminal GST-tagged protein.

The GST-tagged TIL was expressed in BL21-CodonPlus (DE3)-RIL competent cells (Agilent Technologies). Two 1 L cultures were grown from a 50 mL starter culture at 37 °C until the O.D. reached ~0.8. The protein was induced with IPTG (300 µM) and the expression was carried out at 18 °C for 12-16 h. Cells were centrifuged at 5000 rpm for 10 min and the pellets were re-suspended in lysis buffer (50 mM tris, 300 mM NaCl, 1 mM EDTA, 2 mM DTT, pH 8.0), followed by addition of the protease inhibitor phenylmethylsulfonyl fluoride (1 mM). Cells were lysed by tip sonication, and centrifugation at 15000 rpm for 60 min removed the cell debris. The supernatant was filtered through 0.22 µm pore size syringe filters (MilliporeSigma, Millex) and loaded onto a GST trap column (GE healthcare) with the lysis buffer. The bound GST-TIL was washed with the lysis buffer and eluted with a buffer containing 20 mM reduced glutathione along with 50 mM tris, 300 mM NaCl, 1 mM EDTA, 2 mM DTT, pH 8.0. TIL was cleaved from the GST-tag by treating the eluate with PreScission protease (125 µg in 14 mL) at 4 °C for 4-6 h. The protease treated protein mixture was loaded onto a HiTrap SP HP cation exchange column (GE Healthcare) and TIL was eluted by running a gradient of NaCl (buffer A: 20 mM sodium phosphates, 150 mM NaCl, 1 mM EDTA, 1 m TCEP, pH 7.0; buffer B: 20 mM sodium phosphates, 1 M NaCl, 1 mM EDTA, 1 mM TCEP, pH 7.0). Fractions were collected by monitoring the UV absorbance at 254 nm and analyzed via SDS-PAGE. Pure fractions containing TIL were combined and concentrated by Amicon ultra centrifugal filters with 3 kDa molecular weight cutoff (Millipore) and buffer was exchanged to either 20 mM HEPES, 150 mM NaCl, 1 mM TCEP; pH 7.4 (mentioned as HEPES-buffer hereafter) or phosphate buffer (10 mM sodium phosphate, 150 mM NaCl, 1 mM TCEP; pH 7.0) as required.

Molecular weight of the purified peptide was verified through MALDI mass spectrometry: expected m/z 8862 (M+H), observed m/z 8861.5 (Supplementary Figure S1). Purity of the peptide was confirmed with SDS-PAGE (Supplementary Figure S2A). Disordered nature of the peptide was verified by circular dichroism spectroscopy (Supplementary Figure S2B).

**N-terminal labeling of TIL.** TIL was selectively labeled at the N-terminal amine group with Alexa 488-SDP ester. The peptide was buffer-exchanged to phosphate buffer (10 mM sodium phosphates, 150 mM NaCl, 1 mM TCEP; pH 7.0) in order to carry out the labeling reaction. The dye was added at a two-fold molar excess to the peptide and the reaction was carried out at 4 °C for 12-16 h. Progress of the reaction was monitored by MALDI mass spectrometer (m/z 9398 for TIL-Alexa 488) and the reaction was continued until 30-50% labeling of TIL was observed (Supplementary Figure S3). Excess fluorophore was separated from the peptide by passing the reaction mixture through a HiTrap desalting column (GE healthcare), eluted against HEPES buffer. Efficiency of the labeling was determined by measuring the fluorophore concentration (Alexa 488, $\varepsilon_{495}$ 73,000 $M^{-1}cm^{-1}$) via a nanodrop instrument (Thermo Fisher) and the peptide concentration via Bradford method.

**Characterization of TIL by MALDI-TOF-MS.** MALDI mass spectrometry was performed to characterize purified TIL and to monitor progress of fluorophore labeling reactions (Supplementary Figures S1 and S3). About 1 µL of the peptide sample was mixed with 1 µL saturated solution of α-cyano-4-hydroxycinnamic acid (in 1:1 acetonitrile: water with 0.1% trifluoroacetic acid). The mixture was spotted on a MTP 384 ground steel target plate (Bruker Daltonics) and allowed to dry. MALDI spectra were recorded with a Bruker UltrafleXtreme MALDI-TOF-MS (Bruker Daltonics).

**Circular dichroism (CD) spectroscopy.** CD spectra were recorded with a AVIV CD spectrophotometer (Aviv Biomedical, NJ) using a High Precision Quartz Suprasil cuvette (Hellma Analytics) at 25 °C. Typically, 5 µM TIL in 10 mM phosphate buffer containing 150 mM NaCl, pH 7.4 were used for recording CD spectra (Supplementary Figure S2B).

**Purification and labeling of full length endophilin, N-BAR and SH3 domain mutants.** Full length rat endophilin A1 (with C108S, E241C, C294S, C295S mutations), its N-BAR domain (with C108S, E241C mutations) domain were expressed, purified and labeled with Alexa 594-maleimide as described elsewhere [1]. Briefly, proteins were expressed as GST-fusions and purified using GST-affinity chromatography. GST tags were cleaved using PreScission protease and the BAR proteins were further purified by anion exchange and size exclusion chromatography techniques. Labeling at the cysteine residue positions were achieved by incubating the protein with Alexa 594-maleimide overnight at 4 °C.

The SH3 domain of rat endophilin A1 was also expressed as GST-fusion protein using a plasmid generously provided by Volker Haucke's lab. The GST-tag was cleaved using thrombin protease and the SH3 domain was further purified by anion exchange and size exclusion chromatography techniques. The purified SH3 domain was labeled at the N-terminus using Alexa

594-NHS ester following the same protocol used for N-terminal labeling of TIL as described in the previous section.

Protein concentrations were determined by measuring the absorbance at 280 nm ($\varepsilon_{280}$ 17545 $M^{-1}cm^{-1}$) for unlabeled full length endophilin. For all the other cases, the Coomassie (Bradford) Protein Assay (Thermo Scientific, USA) was used for determining protein concentrations.

**GUV preparation and conjugation of TIL to GUVs.** GUVs were prepared by the electroformation method using indium-tin oxide (ITO) coated slides [2]. Chloroform solutions of desired lipid composition were coated onto ITO slides and vacuum dried for at least 2 hours. The lipid films were hydrated with sucrose solution (350 mOsm in MilliQ purified water). Electroformation was performed at 55 ºC for 1 hour.

For TIL conjugation, GUVs composed of mCC-PE/DOPC/DiD (5:94.5:0.5 molar ratio) were mixed (in 1:10 GUV solution: buffer volumetric ratio) with TIL-Alexa 488 (100 nM) in HEPES buffer. The buffer osmolarity was pre-adjusted to the osmolarity of the sucrose solution used for electroformation. The mixture was incubated at room temperature for 6-8 h and then the unreacted maleimide groups on the lipids were blocked by adding β-mercaptoethanol (5 mM).

**Confocal imaging and quantitative image analysis.** Confocal images were recorded using a FluoView 3000 scanning system configured on an IX83 inverted microscope (Olympus, Center Valley, PA). Images were taken at room temperature using a 60 x 1.1 NA water immersion objective (Olympus). GUVs suspended in buffer were placed into the cavity of a home built imaging chamber formed by two glass coverslips (25 × 25 $mm^2$, Fisher Scientific) held together with a glass holder with the help of vacuum grease. The bottom coverslip was passivated with β-casein solution (5 mg/mL) in order to avoid sticking of GUVs to the glass surface. GUVs settled on the bottom glass surface were imaged.

For study of the ionic strength dependent TIL binding to anionic lipid containing membranes, GUVs composed of DOPS/DOPE/DOPC/DiD (45:30:24.5:0.5 molar ratio) were mixed with TIL-Alexa 488 in HEPES buffer containing 0-200 mM NaCl. The osmolarity difference between the inside and outside of the GUVs was balanced by adding requisite amounts of glucose to the outer solution. TIL-Alexa 488 binding to GUVs was determined by imaging via the Alexa 488 channel ($\lambda_{ex}$ 488 nm, $\lambda_{em}$ 500-540 nm).

For the endophilin binding experiments, GUVs conjugated with TIL were mixed with Alexa 594 labeled endophilin before imaging. A three-channel orthogonal imaging sequence was used to image in the Alexa 488 channel, Alexa 594 channel ($\lambda_{ex}$ 560 nm, $\lambda_{em}$ 580-620 nm), and DiD channel ($\lambda_{ex}$ 640 nm, $\lambda_{em}$ 650-750 nm) using three different detectors. The excitation lasers were alternately switched such that crosstalk between different channels could be minimized. Images were analyzed with ImageJ and MATLAB programs. The fluorescence intensity in a given channel was quantified by fitting the GUV contour with a Gaussian ring [3].

**Fluorescence Recovery after Photobleaching (FRAP) experiments.** FRAP experiments were performed with a FluoView 3000 scanning system configured on an Olympus IX83 confocal microscope. TIL-Alexa 488-conjugated GUVs having lipid composition mCC-PE/DOPC/DiD (5:94.5:0.5) were first imaged by excitation with a 488 nm laser at 1% laser attenuation power. Photobleaching was performed at a selected box-shaped area on the GUV surface by applying 488 nm laser at 100% laser attenuation power for 1 second. Images of the recovery stage were collected immediately after the bleaching at a 2.6 seconds per frame image capture speed.

Images were analyzed with ImageJ software. For each time frame, mean intensity of a region of interest (ROI) chosen within the bleached region on the GUV surface was estimated and normalized against the mean intensity obtained from a similar ROI in the unbleached region. The normalized intensities were plotted with respect to time to obtain a FRAP recovery profile (Figure S6B).

**Membrane tether pulling and analysis of curvature sorting.** Membrane tethers were pulled from micropipette aspirated GUVs composed of mCC-PE/DOPC/DiD/DSPE-PEG-Biotin (5:94.3:0.5:0.2 molar ratio) [4]. Glass capillaries (World Precision Instruments (WPI, FL)) were pulled using a pipette puller (Sutter Instruments, CA) and the tips were remodeled with a microforge (WPI). Inner diameters of the capillaries were ~7 µm for GUV aspiration and ~5 µm for capturing beads. The capillary tips were passivated with β-casein solution (5 mg/mL). Capillaries were filled with HEPES buffer using a MicroFil needle (WPI) and fitted to the two arms of a motorized micromanipulator (Luigs & Neumann, Ratingen, Germany) oriented at an angle of 90º. One of these pipettes was used for aspiration of GUVs while a second one held a streptavidin-coated bead to pull tethers from the aspirated GUVs. Aspiration pressure was maintained by adjusting the height of a water reservoir connected to the pipette used for GUV aspiration, and the pressure was monitored by a pressure transducer (Validyne Engineering, Los Angeles, CA). Membrane tension ($T$) was determined from the aspiration pressure ($\Delta P$), radius of the GUV ($R_v$) and the radius of the projection area ($R_p$) by using the formula [5], $T = \Delta P \times \frac{R_p}{2(1-\frac{R_p}{R_v})}$

TIL-conjugated GUVs were mixed with protein (endophilin or N-BAR) and pipette-aspirated. The aspiration pressure was maintained such that the resulting membrane tension was within the range of 0.12±0.2 mN/m. The aspirated GUV was moved upward (by ~ 0.5 mm) from the coverslip surface and incubated until the length of the aspirated vesicle projection stabilized. The tether was pulled by briefly (~1 sec or less) touching the streptavidin bead to the GUV surface and moving it to a distance of 12-15 µm from the surface carefully (at a speed of 2-3 µm/s approximately), in order to ensure that the tether remained intact.

The orientation and length of the pulled tether were adjusted by fine-tuning the x, y, and z positions of the bead with the micromanipulator system such that the tether was oriented along the center axis of the aspiration pipette and showed uniform fluorescence intensity along the tether length. Images were collected in Alexa 488, Alexa 594, and DiD channels. To determine the sorting coefficient, intensities on the flat GUV membrane were determined in MATLAB program as described in previous sections. The aspirated region of the GUV was excluded from the fluorescence intensity analysis. The fluorescence intensity on the tether was quantified in ImageJ software by selecting a narrow, rectangular region of interest (ROI) covering the pixels along the length of the tether and measuring the mean pixel intensity within that ROI.

**LUV preparation and TEM imaging.** LUVs of desired lipid compositions were prepared following a method described elsewhere [6]. Briefly, lipid films were prepared by evaporating chloroform solutions of the desired lipid composition. HEPES buffer was added to the lipid films such that the resulting total lipid concentration was ~1 mM. The aqueous suspension of lipids was vortexed and extruded through 400 nm pore sized polycarbonate membranes (Whatmann/GE Healthcare) in order to obtain LUVs of uniform sizes. Size distribution of the LUVs were confirmed by dynamic light scattering (Malvern, United Kingdom) (Supplementary Figure S4). For the coupling of TIL, LUVs composed of mCC-PE/DOPC (5:95) were incubated (0.1 mM final lipid conc.) with TIL (250 nM) for 8 hours at room temperature in HEPES buffer. The reaction was quenched by adding β-mercaptoethanol (5 mM).

For tubulation assays, LUVs (0.1 mM total lipid conc.) with or without conjugated TIL were incubated with endophilin (5 µM) in HEPES buffer for 30 minutes at room temperature. A droplet (20 µL) of the mixture was added onto a piece of parafilm and a carbon coated copper grid (Electron

Microscopy Sciences, Hatfield, PA) was gently placed on the droplet such that the coated surface faced the liquid in order to allow the vesicles to stick to the grid. The grid was removed after 2 min and excess solutions were soaked with a filter paper (Whatmann). The grid was washed (thrice for TIL conjugated LUVs and once for LUVs having no TIL) by dipping into droplets of HEPES buffer, followed by removal of the excess buffer with a filter paper. For negative staining, the grid was placed on a droplet (20 µL) of 2% uranyl acetate solution for 2 min. Extra stains were washed thrice with buffer and grids were kept on a filter paper for 10 minutes at room temperature for further drying. Images were recorded on a JEM 1011 transmission EM (JEOL, USA), operated at 100 kV, coupled with an ORIUS 832.10 W CCD camera (Gatan). Image analysis and post-processing were performed with ImageJ software.

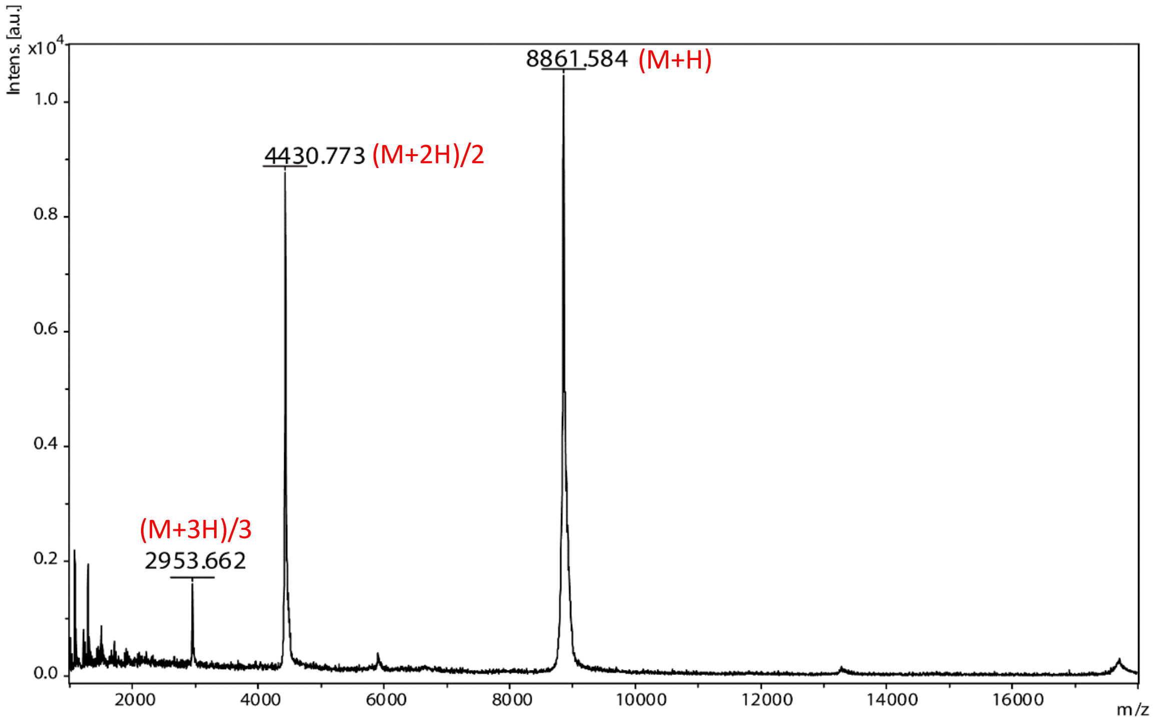


**Figure S1.** Characterization of purified TIL with MALDI mass spectrometry. Expected m/z: 8862 (M+H).

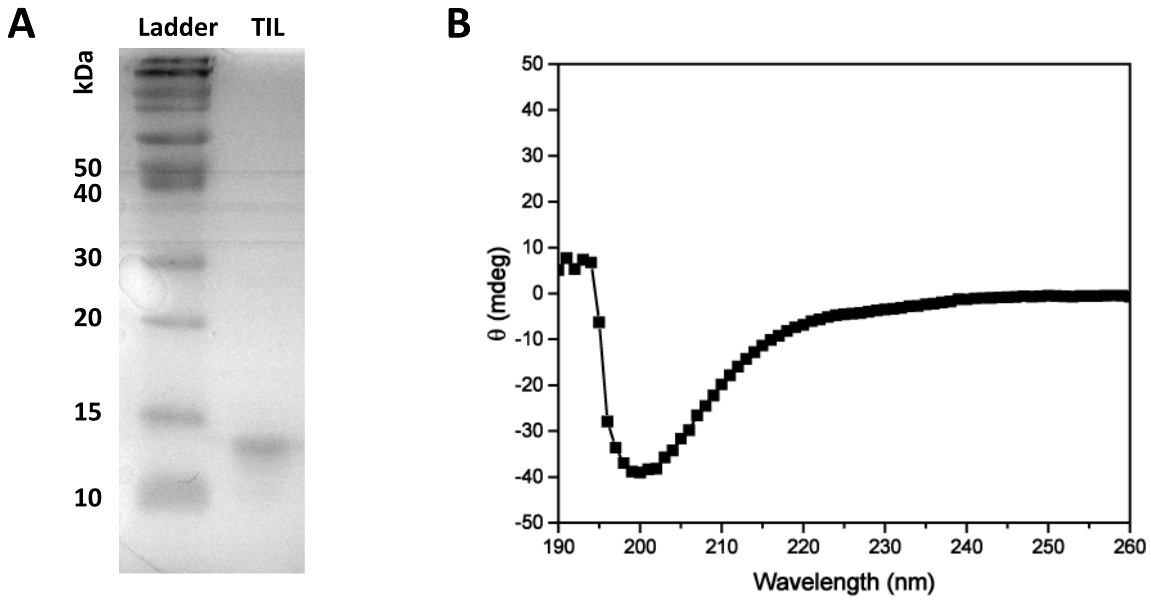


**Figure S2.** A. Characterization of TIL with SDS-PAGE. B. Circular dichroism (CD) spectra of purified TIL.

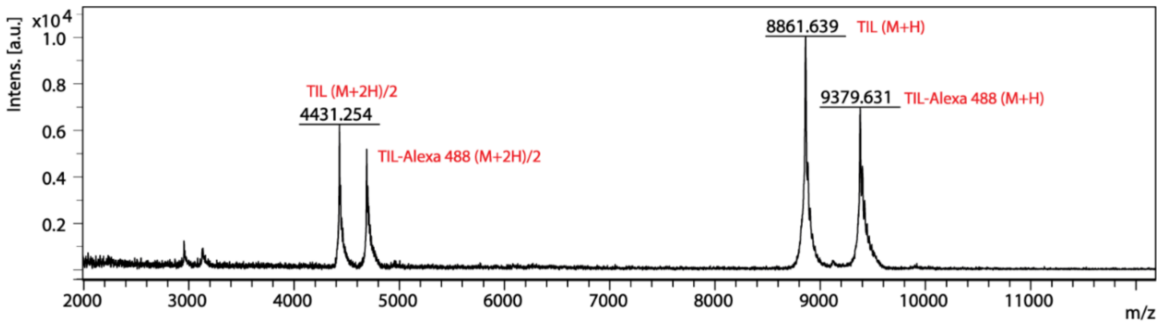


**Figure S3.** MALDI MS spectrum of TIL after labeling with Alexa 488-SDP ester. Expected m/z after labeling: 9379 (M+H).

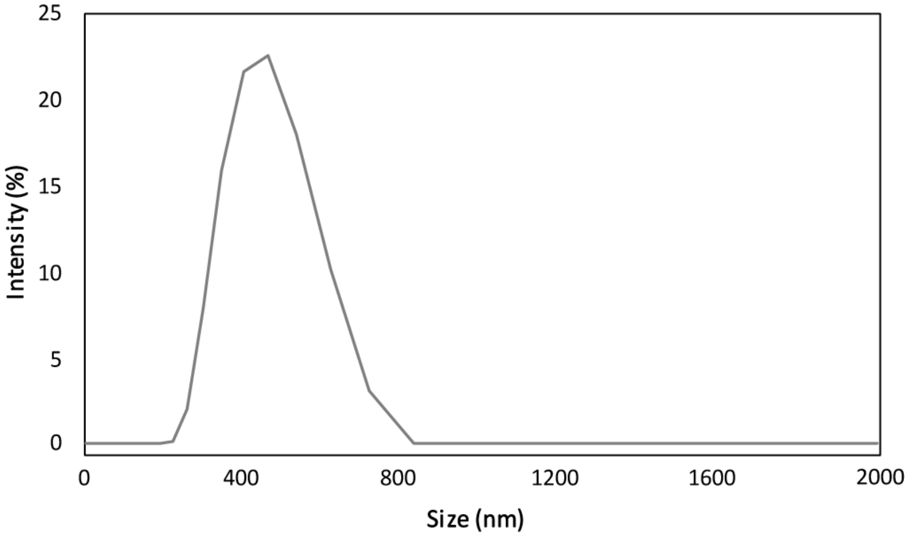


**Figure S4.** Size distribution of LUVs composed of 5% mCC-PE, 95% DOPC as obtained from dynamic light scattering.

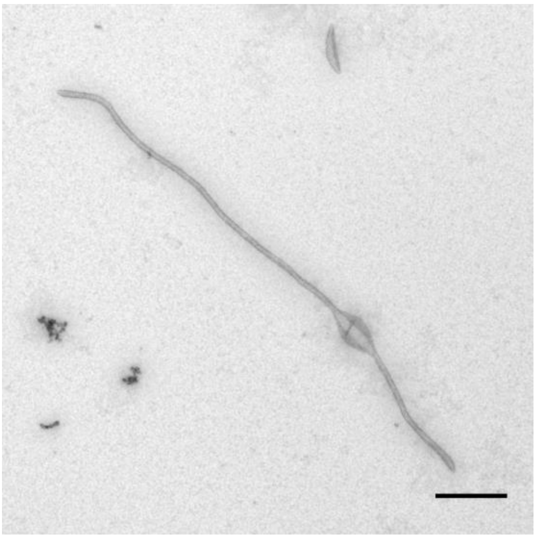

**Figure S5.** TEM image of a tubule formed by PS:PE:PC (45:30:25) LUV in the presence of endophilin (5 µM). Scale bar: 500 nm.

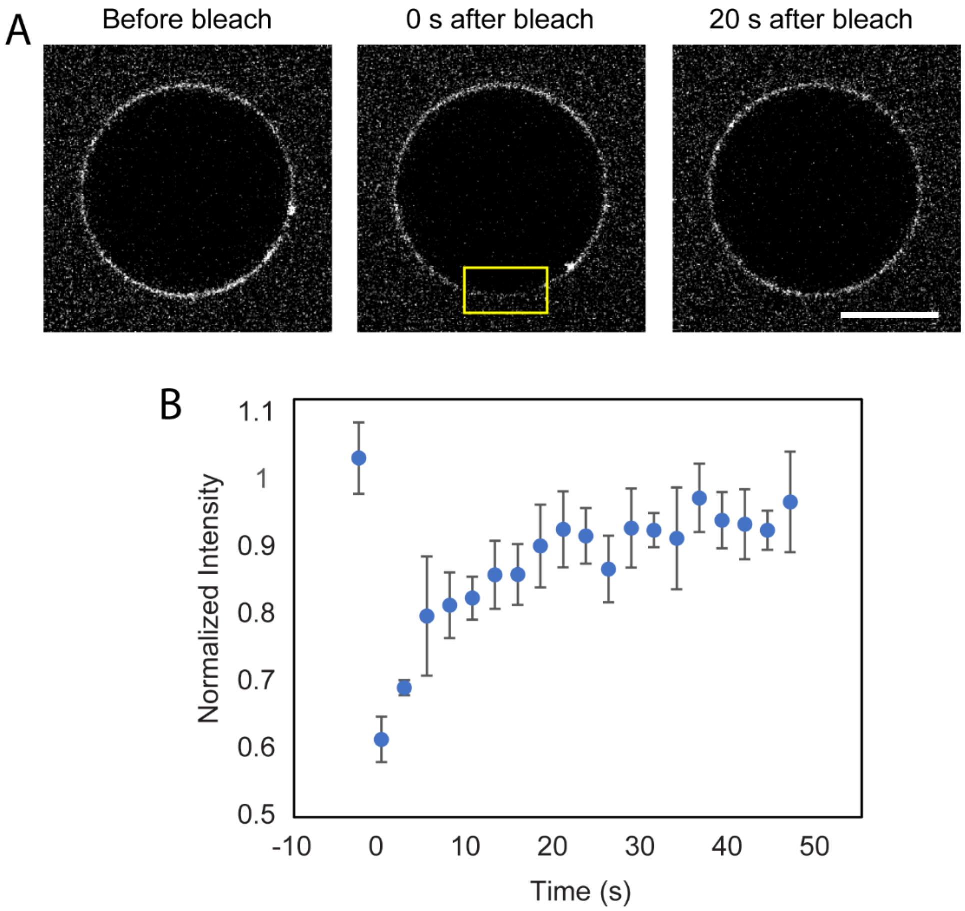


**Figure S6.** Fluorescence recovery after photobleaching (FRAP) studies on TIL-conjugated GUVs. A. Confocal images of TIL-conjugated GUVs (mCC-PE/DOPC/DiD) before, immediately after (0 s time delay) photobleaching, and 20 s after photobleaching. The yellow box indicates the membrane area irradiated for bleaching. B. Recovery profile obtained by plotting the normalized fluorescence intensities from the bleached membrane area with respect to time. Each data point is represented as mean ± standard deviation from five independent FRAP experiments.

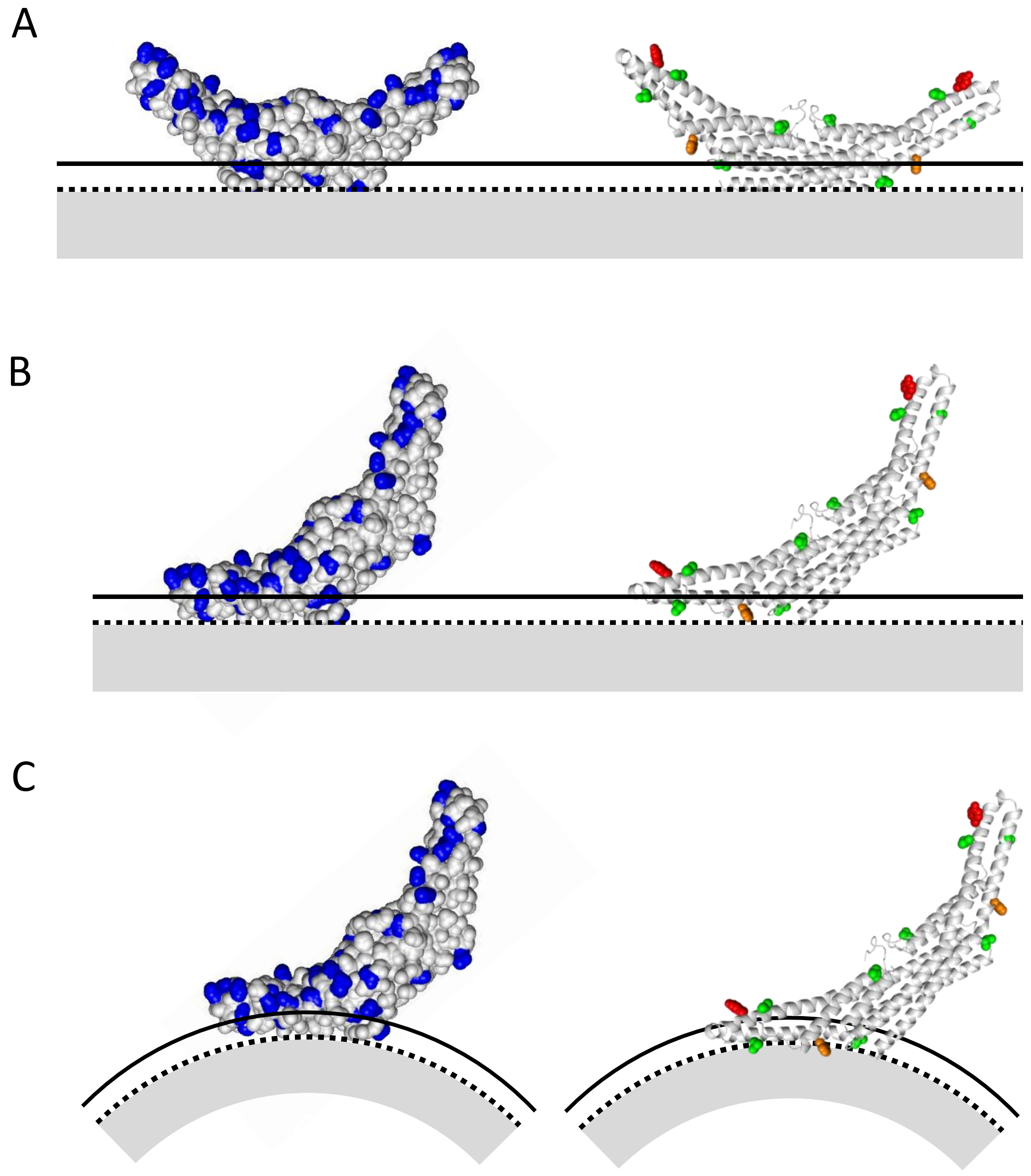


**Figure S7:** Membrane interaction of hydrophobic and cationic sidechains at various orientations of the BAR domain with respect to the membrane. A. BAR domain facing flat membrane with the top of the convex surface. Surface accessible Lys and Arg residues (blue) that can interact with phosphate group of the PC lipids are shown on the left. On the right, positions of the surface accessible hydrophobic residues (green: aliphatic, red: aromatic) are shown for the same orientation. Solid line represents the top of the choline headgroups, dotted line shows region containing phosphates and the grey indicates lipid tail region. B. Membrane interactions of cationic (left) and hydrophobic (right) residues when the protein is facing the membrane with one side of the convex surface. C. Membrane interactions of cationic (left) and hydrophobic (right) residues on a curved membrane when the protein is facing the membrane with one side of the convex surface.

**SI References**


1. Capraro, B. R.; Shi, Z.; Wu, T.; Chen, Z.; Dunn, J. M.; Rhoades, E.; Baumgart, T., Kinetics of endophilin N-BAR domain dimerization and membrane interactions. *The Journal of biological chemistry* **2013,** *288* (18), 12533-43.
2. Mathivet, L.; Cribier, S.; Devaux, P. F., Shape change and physical properties of giant phospholipid vesicles prepared in the presence of an AC electric field. *Biophys J* **1996,** *70* (3), 1112-21.
3. Shi, Z.; Baumgart, T., Membrane tension and peripheral protein density mediate membrane shape transitions. *Nat Commun* **2015,** *6*, 5974.
4. (a) Sorre, B.; Callan-Jones, A.; Manneville, J. B.; Nassoy, P.; Joanny, J. F.; Prost, J.; Goud, B.; Bassereau, P., Curvature-driven lipid sorting needs proximity to a demixing point and is aided by proteins. *Proc Natl Acad Sci U S A* **2009,** *106* (14), 5622-6; (b) Tian, A.; Baumgart, T., Sorting of lipids and proteins in membrane curvature gradients. *Biophys J* **2009,** *96* (7), 2676-88.
5. Waugh, R.; Evans, E. A., Thermoelasticity of red blood cell membrane. *Biophys J* **1979,** *26* (1), 115-31.
6. Chen, Z.; Zhu, C.; Kuo, C. J.; Robustelli, J.; Baumgart, T., The N-Terminal Amphipathic Helix of Endophilin Does Not Contribute to Its Molecular Curvature Generation Capacity. *J Am Chem Soc* **2016,** *138* (44), 14616-14622.
